# Assessment of binocular fixational eye movements including cyclotorsion with split-field binocular scanning laser ophthalmoscopy

**DOI:** 10.1101/2022.03.03.482756

**Authors:** Julia Hofmann, Lennart Domdei, Stephanie Jainta, Wolf Harmening

## Abstract

Fixational eye movements are a hallmark of human gaze behavior, yet little is known about how they interact between fellow eyes. Here, we designed, built and validated a split-field binocular scanning laser ophthalmoscope (bSLO) to record high-resolution eye motion traces from both eyes of six observers during fixation at different binocular vergence conditions. In addition to microsaccades and drift, torsional eye motion could be extracted, with a spatial measurement error of less than 1 arcmin. Microsaccades were strongly coupled between fellow eyes under all conditions. No monocular microsaccade occurred and no significant delay between microsaccade onsets across fellow eyes could be detected. Cyclotorsion was also firmly coupled between both eyes, occurring typically in conjugacy, with gradual changes during drift and abrupt changes during saccades.

## Introduction

Seeing with two eyes, i.e. binocularity, is central to human visual processing. For instance, ocular controls of the retinal image forming process, like pupil constriction and accommodation, are highly coupled between the two eyes (Flitcroft, Judge & Morley, 1992), and ocular motor commands issued to control gaze of one eye are tightly coupled to those of the fellow eye (Tweed, 1997; Murray, Gupta, Dulaney, Garg, Shaikh & Ghasia, 2022), for example during tracking of a moving object. Less is known whether this tight coupling extends to the phases of stable fixation, where fixational eye movements (FEM) predominate (Krauskopf, Cornsweet & Riggs, 1960; Otero-Millan, Macknik & Martinez-Conde, 2014; Simon, Schulz, Rassow & Haase, 1984). For example, Krauskopf, Cornsweet, and Riggs (1960) emphasized that microsaccades occur synchronously in both eyes (Krauskopf, Cornsweet & Riggs, 1960), but this early finding is still discussed controversially (Møller, Laursen, Tygesen & Sjølie, 2002; Engbert & Kliegl, 2003; Zhou & King, 1998). Binocular eye movements in general cannot be reduced to the yoking of the eyes during saccades: vergence eye movements occur as horizontal, vertical or cyclovergence. All three movements show substantial differences in their contributions to fusion (i.e the perception of a single image; Leigh & Zee, 2006; Schor & Ciuffreda, 1983; Steinman, Steinman, & Garzia, 2000): while horizontal vergence, for example, reacts to – on large scale and fine-tuned – horizontal disparity of the object that needs to be foveated, vertical vergence reacts to vertical misalignments of the whole image of one eye relative to the other eye. Vertical eye movements are supposed to be inherently conjugate in that vertical premotor neurons simultaneously drive both eyes (McCrea, Strassman & Highstein, 1987). Torsional eye movements (cyclovergence) are also small in overall variability (about 0.10°), and are thus tightly controlled (Van Rijn, van der Steen & Collewijn, 1994). According to van Rijn et al. (1992) cyclovergence is a truly binocular process and, unlike cycloversion, requires correspondence of the images presented to the two eyes. Finally, tremor represents a small periodic eye movement (Martinez-Conde et al., 2004; Rolfs, 2009), but whether tremor is a binocularly coordinated eye movement is still discussed (Riggs & Ratliff, 1951; Spauschus et al., 1999). Very few data exist showing all binocular eye movements – systematically – during FEM.

FEM are very small in amplitude, typically just a few minutes of arc of visual angle, and are thus not trivial to observe and to measure accurately (Rucci & Victor, 2015; Rolfs,2009). To study FEM, both spatial and temporal resolution of the measurement technique needs to be high (Poletti & Rucci, 2015; Otero-Millan, Macknik, Langston & Martinez-Conde, 2013; Chung, Kumar, Li & Levi, 2015). Such techniques include invasive means by attaching mirrors or coils directly to the moving eyeball (Barlow, 1952), or non-invasive means such as pupil and Purkinje image video-tracking (Poletti & Rucci, 2015; Martinez-Conde, Otero-Millan & Macnik, 2013), and retinal tracking by scanning laser ophthalmoscopy (SLO) (Stevenson, Roorda & Kumar, 2010; Sheehy, Yang, Arathorn, Tiruveedhula, Boer & Roorda, 2012). Owing to the obvious disadvantages of invasive measurement techniques, the highest spatial precision is currently achieved by SLO (Sheehy, Yang, Arathorn, Tiruveedhula, Boer & Roorda, 2012). If combined with potent image registration tools and micro-stimulation tools, SLO-based retinal tracking can serve as a highly sensitive gaze tracker with minimal spatial distortions artifacts (Bowers, Boehm & Roorda, 2019), allowing also to directly observe the retinal location of fixation of a visual target (Stevenson, Roorda & Kumar, 2010; Vogel, Arathorn, Roorda & Parker, 2006).

In this work, we describe an improved binocular scanning laser ophthalmoscope with which fixational eye motion can be studied during binocular vision with relative ease. We measured binocular fixational eye movements in six healthy participants. Next to high-resolution measurements of binocular gaze behavior, our analysis also allowed to extract cyclotorsion, the rotation of the eyeballs around the visual axes, with high spatial resolution.

## Methods

### Binocular scanning laser ophthalmoscope, bSLO

A binocular scanning laser ophthalmoscope (bSLO) was developed, similar to an earlier design described by Stevenson, Sheehy and Roorda (2016), with additional functional improvements (see **Fig. 1**). Pertinent details are described here. A mirror-based (f = 300 mm) SLO with confocal detection scheme was designed in optical simulation software (Zemax Optics Studio, Zemax Germany GmbH, Munich, Germany), and optimized to allow diffraction limited lateral resolution across a 3×3 deg field of view in each eye (see spot diagrams in **Fig. 1D**). Beam folding of the afocal, 4-f telescopic front-end followed orthogonal folding rules as laid out in Gomez-Vieyra et al. (2009) to minimize system astigmatism. Light source was the fiber-coupled output of a superluminescent light emitting diode with 795 nm center wavelength (∼15 nm FWHM) (SLD-CS-381-HP3-SM-795-I, Superlum, Cork, Ireland). After launching a 3.6 mm-diameter collimated beam in the reflection portion of the a 50:50 beam splitter into the bSLO front-end, galvanometric (30 Hz sawtooth) and resonant (∼16 kHz sinusoidal) scanning, positioned in conjugate pupil planes, produced a raster field size of 3 × 6 (vertical x horizontal) degrees of visual angle. A knife-edge mirror (Thorlabs MRAK25-P01), placed in a retinal plane, split the rectangular raster into two square, 3 × 3-degree half-fields, which were optically relayed separately into both eyes via a lens-based Badal optometer (Range of correctible ocular defocus: +2 to -7D). The last fold mirrors before the eyes were on single-axis translation and rotational stages that could be operated electronically to correct for interpupillary distance and binocular vergence angle. Power incident at each cornea was 200 µW. The light reflected by the retina was detected in the transmitted portion of the 50:50 beam splitter in a single photomultiplier tube (H7422-50, Hamamatsu Photonics, Hamamatsu, Japan), placed behind a confocal pinhole (pinhole diameter = 50 µm, equaling 0.9 Airy disk diameters). PMT output signals were sampled at 20 MHz by a field programmable gate array (FPGA) in custom software (ICANDI, available at https://github.com/C-RITE) to produce 512×512 pixel video frames at 29.3 Hz.

**Figure 1.**
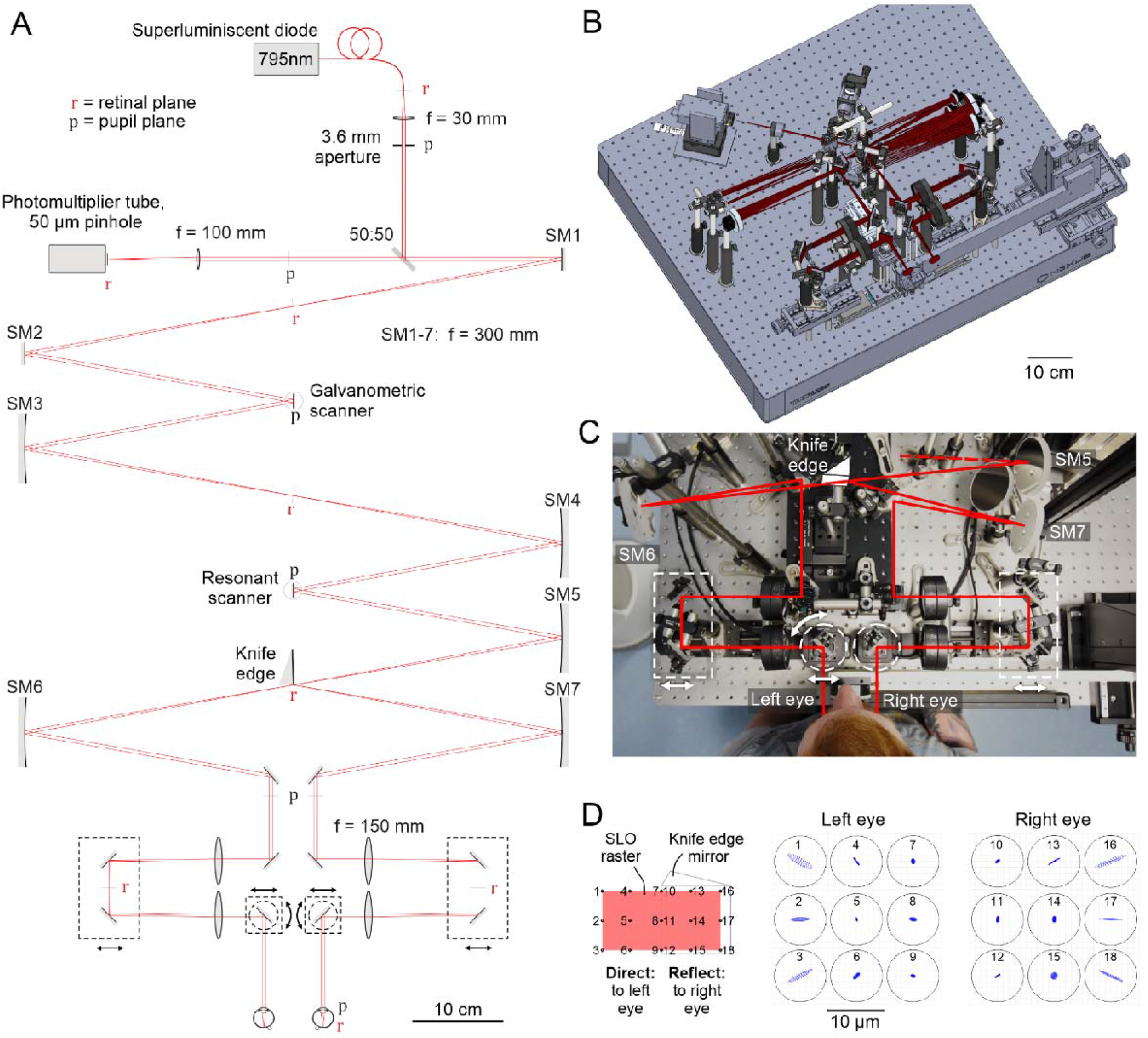
Binocular scanning laser ophthalmoscope (bSLO). **A**: Schematic drawing of the bSLO setup. Components are drawn to scale; the beam path is shown unfolded for clarity (compare **B** for actual beam path). Translation and rotation stages marked by dashed lines. Scale is given along the direction of beam propagation. **B**: Three-dimensional model of the actual beam path and optomechanical components. **C**: Top view photograph of the front-end with indication of beam paths for both eyes and position of moveable stages. **D**: Simulated spot diagrams at the 18 cardinal points of the bSLO raster, spanning square imaging fields in the two eyes. Circles indicate Airy disk diameter.

### Eye motion extraction from bSLO videos

Due to the field-split design of the bSLO, a single video frame consisted of two half images of each retina recorded side-by-side (**Fig. 2A**). Because of the equal aspect ratio in the FPGA digital sampling and the rectangular optical scanning field with an aspect ratio of 1:2 (horizontal:vertical), retinal image space in each half-image was compressed along the horizontal dimension two-fold. Digital resolution was 84 pixels/degree in the horizontal direction and 168 pixels/degree in the vertical direction. This anisotropy was compensated later by multiplying horizontal motion signals by 2. Binocular eye motion extraction was achieved by an offline strip-wise image registration described earlier of each half-field independently (Stevenson, Roorda & Kumar, 2010). In brief, half images were divided into 32 horizontal strips, each 16 pixels high, and registered to a high-definition reference frame, generated automatically from a longer video sequence (**Fig. 2B**). This produced a high-resolution eye motion trace with 960 Hz temporal sampling frequency. From each video, horizontal and vertical motion traces of both eyes were further analyzed. In those traces, microsaccades were semi-manually labelled by first thresholding motion velocity, and then validating each candidate saccade manually (**Fig. 2C**). By setting a velocity threshold at 0.25 arcmin/ms in a moving average of 7 positional samples, a candidate microsaccade was detected. The precise temporal onset of such candidate was then found at the first sample exceeding a velocity of 0.25 arcmin/ms in a moving average of 3 data samples within a 1 frame window around this sample. Microsaccade offset was determined similar to onset, at the first sample where positional velocity dropped below 0.25 arcmin/ms in a moving average of 3 data samples after onset. All candidate microsaccades were manually validated. In the horizontal direction, eye movements shifting gaze to the right were expressed by positive value changes. In the vertical, positive value changes mean gaze upwards (both directions in the visual field as seen from behind the participant).

**Figure 2.**
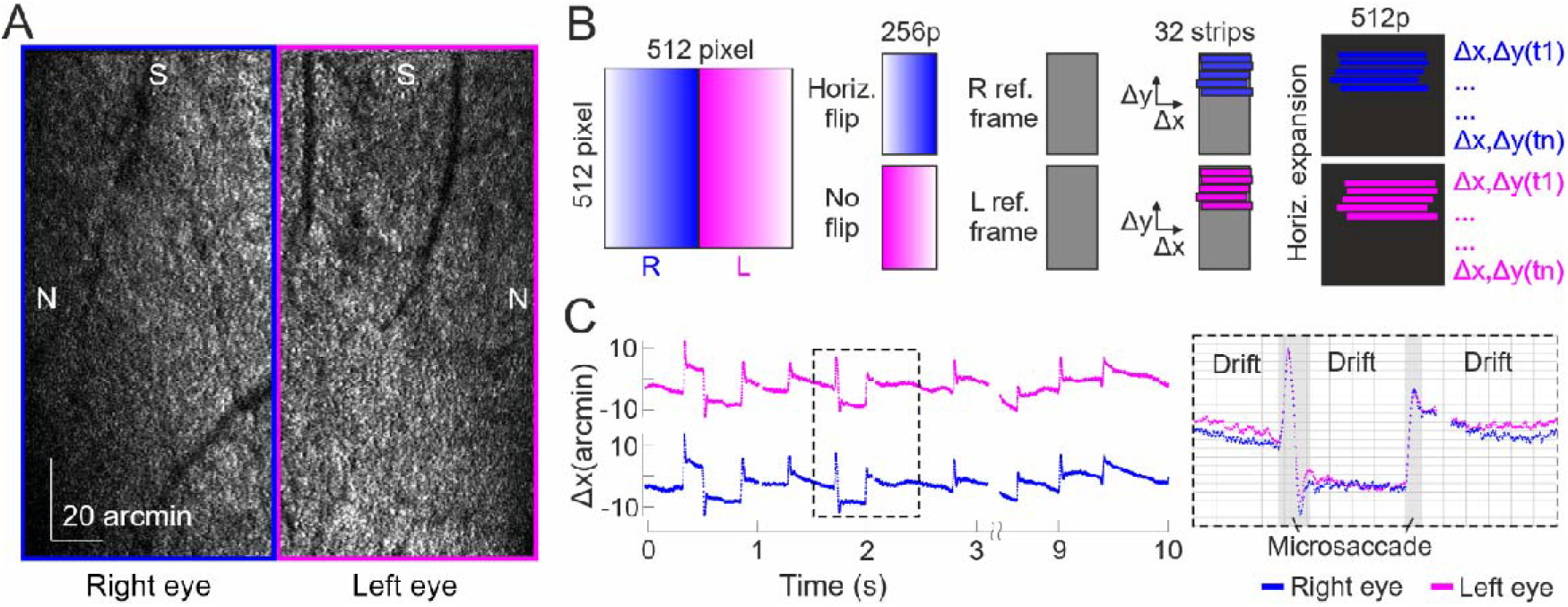
Eye motion extraction from bSLO imagery. **A:** Single bSLO video frame while the participant fixated on the upper right corner of the scanning raster in both eyes. Note that the right eye’s image is horizontally flipped due to the scan geometry and beam folding (S: superior, N: nasal on the retina). **B**: Half-images in single bSLO frames were separated and brought into fundus orientation before strip-wise image registration. Horizontal image shifts were multiplied by 2 to account for the unequal image aspect ratio. **C:** Example retinal motion traces of left and right eyes (horizontal motion only).

### Estimation of cyclotorsional eye motion

Positional eye motion traces resulted from strip-wise image translations relative to a reference image. If the acquired image is however rotated against that reference, e.g. during cyclotorsional movement of the eye, eye motion traces will contain an additional horizontal component beating at frame rate, resembling a sawtooth pattern. This particular rotational artifact is pronounced in the horizontal dimension due to the predominantly horizontal geometry of the image strips (**Fig. 3A**). By computationally rotating the reference image prior to the strip-wise image correlation systematically, we found a linear relationship between the slope at which horizontal strip offsets appeared at frame rate. We selected a total of 604 bSLO video frames from different viewing conditions and eyes where no rotation artifact was visible. Reference images were rotated within the interval 0 to 30 arcmin. From this, we derived a factor of 3.77 between the measured gradient in horizontal positional motion traces (in arcmin per frame) and angle of image rotation (in arcmin) (**Fig. 3B**). The slope of the horizontal motion trace was measured frame-wise using a linear fit to all samples within one frame and converted to image rotation by the aforementioned factor. Cyclotorsion signals could thus be derived at frame rate in all bSLO eye motion traces. Due to the large change in slope during a microsaccade, these epochs were excluded from cyclotorsional analysis (**Fig. 3C**). Motion trace slope that was due to simultaneous drift was separated from torsion signals. For this, the drift slope was calculated by the difference of the mean drift per frame and then subtracted from the torsion slope, leaving the isolated torsion value. Throughout this paper, positive torsional values correspond to clockwise eye rotation as seen from behind the participant.

**Figure 3.**
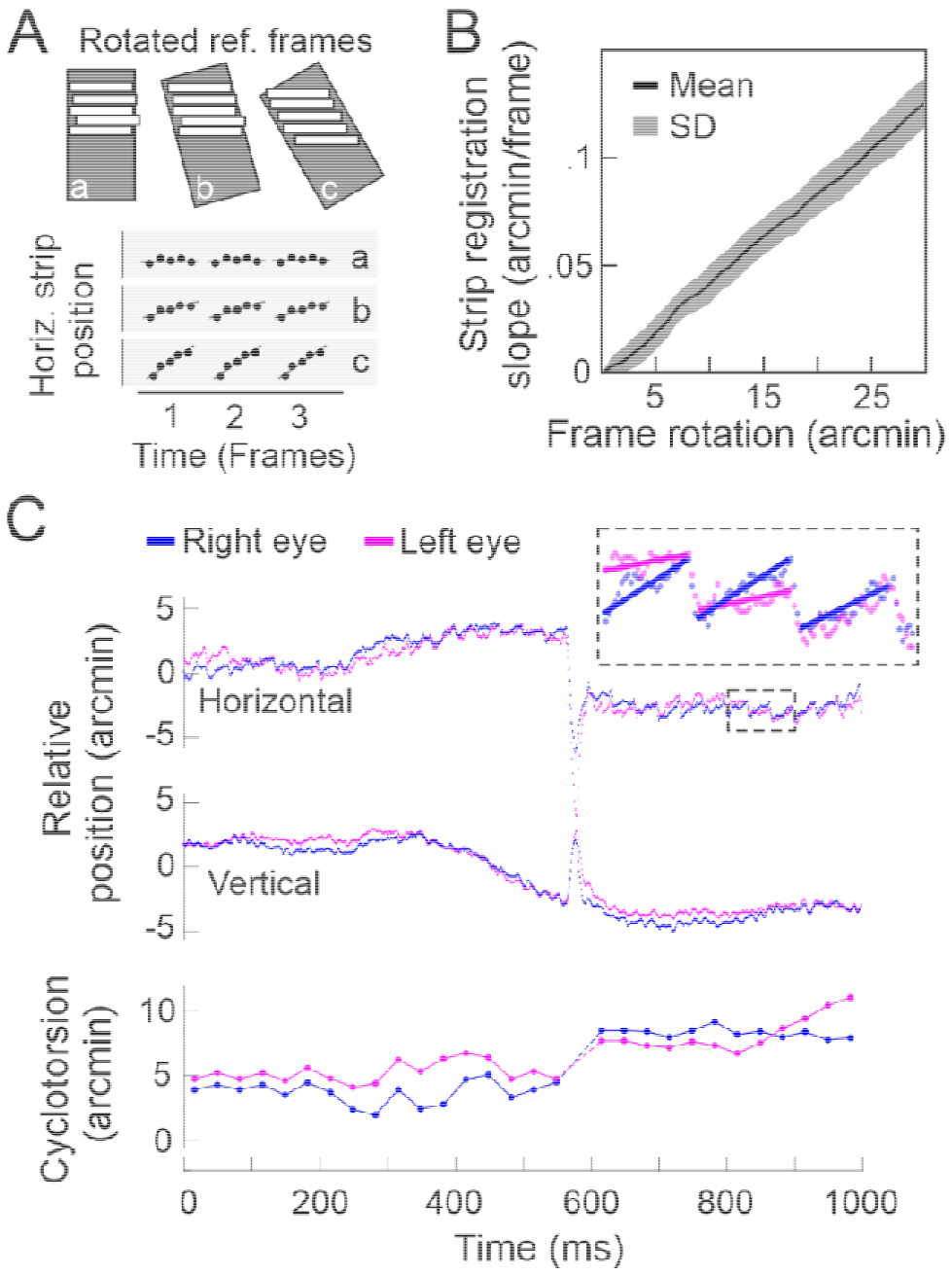
Estimation of cyclotorsional eye motion. **A:** Strip-wise image correlation to a rotated reference frame produces a sawtooth pattern in the horizontal position signal, with the slope being a function of frame rotation. **B:** By computationally rotating the reference frame in a number of image sequences that contained no sawtooth pattern, a linear relationship between strip gradient and rotation angle was determined per frame. **C:** Exemplary torsional analysis with the sawtooth pattern highlighted in the inset. Typically, torsion signals changed with a microsaccade.

### Assessment of binocular FEM

Binocular fixational eye movements (bFEM) were measured in six healthy volunteers (one female, five males, mean age: 34), referred to as P1 to P6 throughout the manuscript. Participant naming was based on a decreasing order of the magnitude of fixational stability, with P1 exhibiting the lowest average binocular deviance iso-contour areas (see **Results**). Refractive state was measured by an auto-refractor and was between -0.125 and - 3 diopters best spherical equivalent. While the participant’s head was immobilized in front of the bSLO by a dental impression (bite bar) held on a XYZ-translation stage, they were asked to fixate on the top right corner of the individual scan raster seen by each eye as relaxed and accurate as possible. To facilitate observer alignment in front of the system, the transversal position of the last fold mirrors was adjusted to accommodate interpupillary distance (IPD). IPD adjustment and observer head positioning followed a simple protocol. First, observer IPD was measured with a handheld digital pupillometer. This reading was entered into a custom written software that controlled the movable stages electronically, and the last fold mirrors travelled to the prescribed distance, symmetrically about the systems center. When the observer then sat in front of the system, only minor misalignments remained, which could be corrected promptly. First, a possible vertical asymmetry of the observer pupil position relative to the parallel system beams was corrected by rotating the gimbal mount which held the bite bar. This head rotation was only necessary for one of the participants (at 2 degrees). In this case, the optimal rotation angle could be found by observing relative bSLO image brightness while the head was moved along the vertical direction via the x,y,z-stage. If the two half images reached maximum brightness at different heights (e.g. right eye lower), the gimbal had to be rotated accordingly (right eye down). A remaining small horizontal asymmetry in pupil position was more common and easily corrected by translation of the whole head relative to the two beams via the x,y,z-stage. Binocular vergence of the last fold mirror of the bSLO was set to either 0, 1, 2, 3, 4 or 5 degrees for each video, in ascending or descending order for half of the subjects, respectively. This was done to both test feasibility of such experimental option and to put a vergence load onto the motor system to trigger differences in FEM dynamics. Five ∼10-second long bSLO videos were recorded at each viewing condition (one video comprised 300 frames = 10.24 s). Pupils were dilated by instilling one drop of 1 % Tropicamide 15 minutes before the beginning of the recording session. Written informed consent was obtained from each participant and all experimental procedures adhered to the tenets of the Declaration of Helsinki, in accordance with the guidelines of the independent ethics committee of the medical faculty at the Rheinische Friedrich-Wilhelms-Universität of Bonn, Germany.

## Results

### Binocular coordination of fixational eye movements

In all six participants (P1-P6), binocular FEM were derived from thirty ∼10-second videos during six different binocular vergence angles (5 in each condition). In each video, horizontal and vertical movements of both eyes were extracted at 960 Hz, torsion was extracted at 30 Hz (see Methods). After removing video frames containing eye blinks and frames that could not be registered to the reference frame (due to out of field motion and other registration errors), we arrived at a total of 1,564,640 samples collected for transversal movement for all eyes combined, and after an additional exclusion of microsaccade epochs, at a total of 37,228 samples for torsional movement. Across all eyes and conditions, positional resolution was high. The variance between adjacent positional samples around a moving average of 10 samples was a tenth of an image pixel, equaling 0.07 arcmin in x-direction and 0.04 arcmin in y-direction. Relative vergence and version were computed by subtracting or averaging the left and right eye motion traces, respectively (an example data set is shown in **Fig. 4** of P4 at a vergence angle of 2° degree).

**Figure 4.**
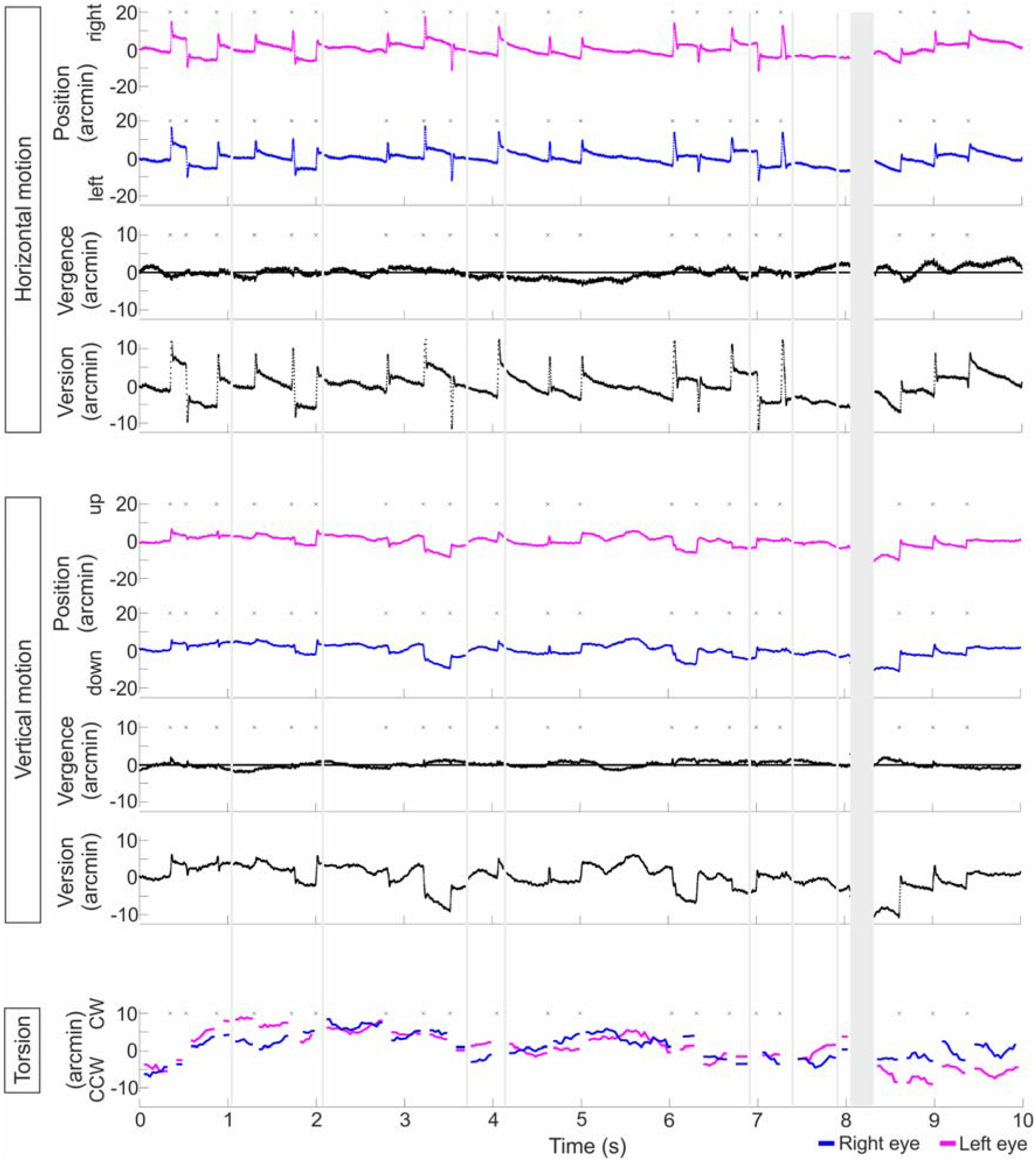
Exemplary binocular eye motion traces. Data is from a single 10-second video recorded in P6 at two-degree vergence angle. Ten of such videos were recorded per viewing condition per participant. Colors indicate fellow eyes (left = magenta, right = blue). Horizontal, vertical and torsional movements are shown separately. Note that for horizontal and vertical movements, positional distances are reported (in arcmin), while for torsion, angular rotation is reported (also in arcmin). Small asterisks indicate the occurrence of a microsaccade for which torsion was undefined.

In general, binocular eye coordination was high for all subjects across all conditions. The average vergence motion (computed as L-R position signals) was 1.18 arcmin (standard deviation, SD: 1.21 arcmin) in the horizontal, and 0.56 arcmin (SD: 0.53 arcmin) in the vertical direction. Within 4,052 total microsaccades detected across all subjects, we did not observe a monocular microsaccade, i.e. one that was present only in one eye. Microsaccade frequency varied across participants (average microsaccades per second, P1: 1.12, P2: 0.71, P3: 0.62, P4: 1.68, P5: 1.41). Temporal microsaccade onset difference between eyes was distributed normally around an average of 1.03 ms (SD: 1.23 ms). The main sequence of microsaccades, defined as the relationship between peak velocity and excursion amplitude showed a typical linear relationship in log-log plotting, with an average slope of 0.073 ms across eyes, equal for fellow eyes (Pearson’s correlation between the left and right eye >0.99 for all subjects). Microsaccade amplitude and direction were firmly coupled between the two eyes (**Fig.5**). Microsaccade amplitude range across all participants was 2.04 to 31.3 arcmin, median amplitude was 13.46 arcmin (N= 4052). The amplitude deviance between left and right (L-R) had a mean of -0.2 arcmin and standard deviation of 1.7 arcmin (Range: 0 – 12.74 arcmin). The mean polar direction deviance was 0.39 degrees, with a standard deviation of 3.68 deg, and a range of 1.32 arcmin to 10.15 deg. Drift amplitudes ranged from 0 to 7.15 arcmin, median amplitude was 2.16 arcmin (N= 7244). Here, the largest absolute amplitude deviance between left and right was 5.8 arcmin (mean: 0.83 arcmin, SD: 0.77 arcmin). The mean direction deviance was 0.54 degrees, with a standard deviation of 34.65 deg, and a range of 0 arcmin to 97.82 deg.

**Figure 5.**
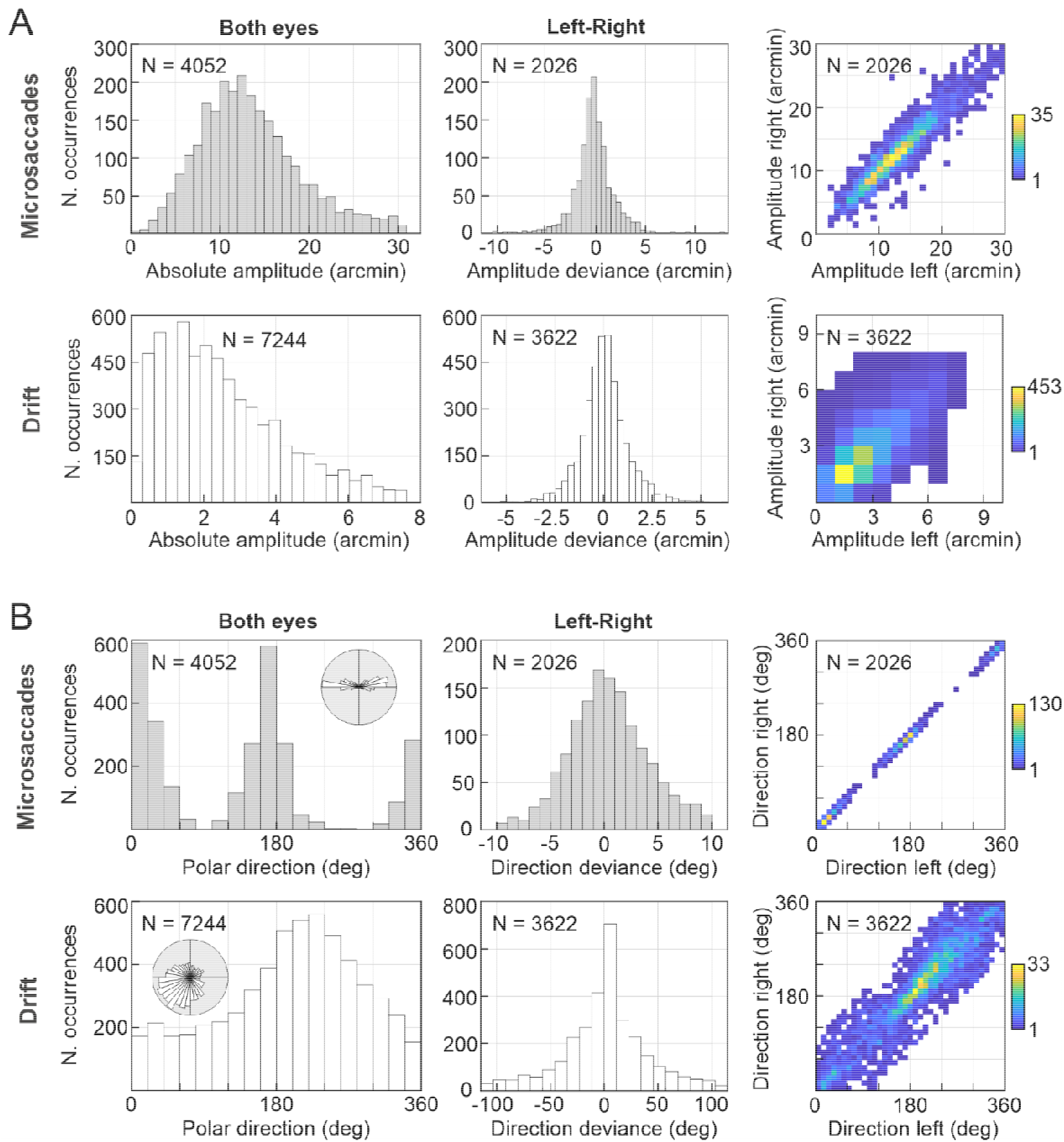
Binocular coupling of microsaccades and drift. A: Analysis of microsaccade and drift amplitude. First column is the histogram of all absolute motion amplitudes, middle column is the histogram of left and right eye amplitude deviances (computed by left-right amplitudes), and third column is the amplitude correlation between all fellow eyes. B: Analysis of microsaccade and drift direction in the visual field (0 deg = right, 90 deg = up). The columns are the same as in A. The small inset in the direction histogram show the same data in polar coordinates for reference.

Across all eyes and viewing conditions, more horizontally oriented microsaccades were performed. Drift direction was mainly pointing down and left, a bias likely induced by the positioning of the fixation target at the upper right corner of the imaging raster.

In an analysis of fixation stability, expressing all retinal landing points of the fixated object in a two-dimensional plot as their iso-contour area (ISOA, encompassing 68% of all data points), a corresponding relationship emerged (**Fig. 6**). While the magnitude of monocular ISOAs differed between participants (average ISOA: P1: 15.62 arcmin^2^, P2: 24.97 arcmin^2^, P3: 27.04 arcmin^2^, P4: 28.46 arcmin^2^, P5: 29.57 arcmin^2^, P6: 31.02 arcmin^2^), fellow-eye ISOAs were similar (Pearson’s correlation between ISOAs of the left and right eye for each participant: P1: ρ=0.78, P2: ρ=0.84, P3: ρ=0.79, P4: ρ=0.62, P5: ρ=0.61, P6: ρ=0.96, all p<<0.01). Binocular deviance ISOAs (L-R) were always smaller than monocular ISOAs (average: 3.48 to 8.04 arcmin^2^, P1 to P6, respectively, average range of monoISOA:binoISOA = 4.52 at binocular vergence of 0°). Binocular deviance ISOAs, unlike monocular ISOAs, were elongated in the horizontal direction, being on average 2.2 times wider than high. We observed a weak yet statistically insignificant trend of increasing monocular ISOAs with larger vergence angles set in the bSLO. However, binocular coupling did not seem to be systematically disturbed by the vergence induced. In all participants, and across all vergence angles, binocular fixation stability (L-R) was always lower then monocular fixation stability (average ISOA ratio mono/bino deviance, 0°: 4.98, 1°: 7.16, 2°: 8.29, 3°: 8.78, 4°: 7.13, 5°: 7.93).

**Figure 6.**
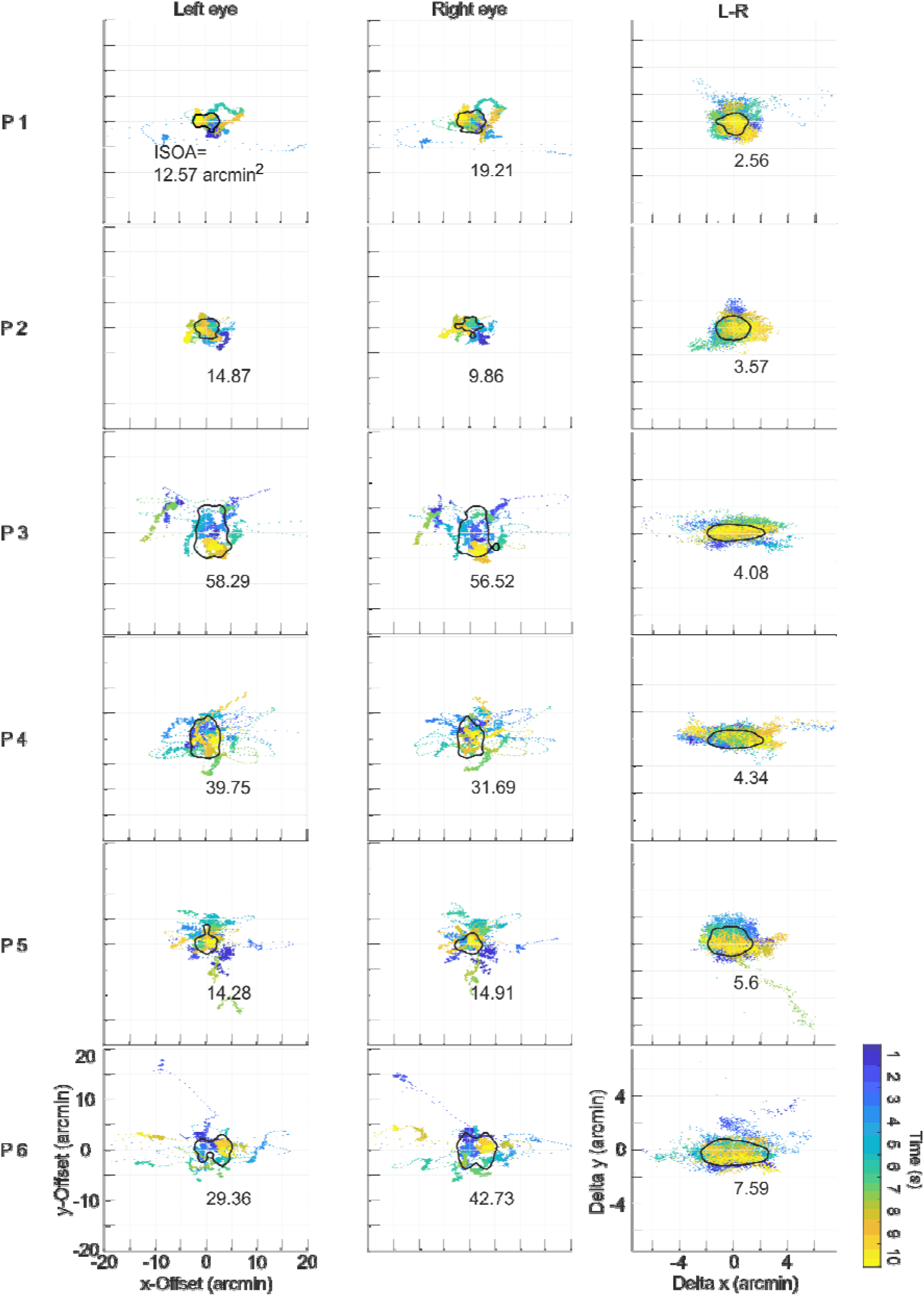
Monocular and binocular fixational stability. Data is from one 10-second video at 0 deg vergence angle for all participants (P1-P6). Fixation target was the corner of the 3×3 degree scanning raster. Time is color coded. The ISOAs (black outline) are given in arcmin^2^. Note that the scale has been magnified for the binocular deviance (L-R) data set 2.5-fold.

### Cyclotorsion during fixational eye motion

From the computationally rotated reference frame analysis we could derive a variance of strip offset for each rotational angle. The smallest strip offset which could be observed had a spatial distance of 0.7 arcmin, derived from the smallest possible slope of the sawtooth pattern measured. The smallest torsional signal which could be measured had a rotational angle of 0.6 arcmin. The square root of the average angular variance (SD: 0.47 arcmin) was multiplied by 2.77 to arrive at a repeatability of 1.3 arcmin. Measurement error was thus 0.92 arcmin (variance multiplied by 1.96). Across all eyes and viewing conditions, torsional angles between -22.9 and 21.6 arcmin were observed (average: 0.53 arcmin, SD: 5.36 arcmin). The largest absolute amplitude deviance between left and right was 11.5 arcmin (mean: 1.57 arcmin, SD: 1.44 arcmin) (**Fig. 7 A)**. Out of all torsion signals (N= 74,456), 36,202 were in counterclockwise direction while 38,254 where in clockwise direction. A linear fit to a correlation of left and right eyes’ torsion demonstrated tight coupling (mean slope = 1.05, sigma^2^ = 0.0005) (**Fig. 7B**). Similar to the metrics of fixation stability and frequency of microsaccades, the distribution of torsional motion was idiosyncratic across participants (**Fig. 7C**).

**Figure 7.**
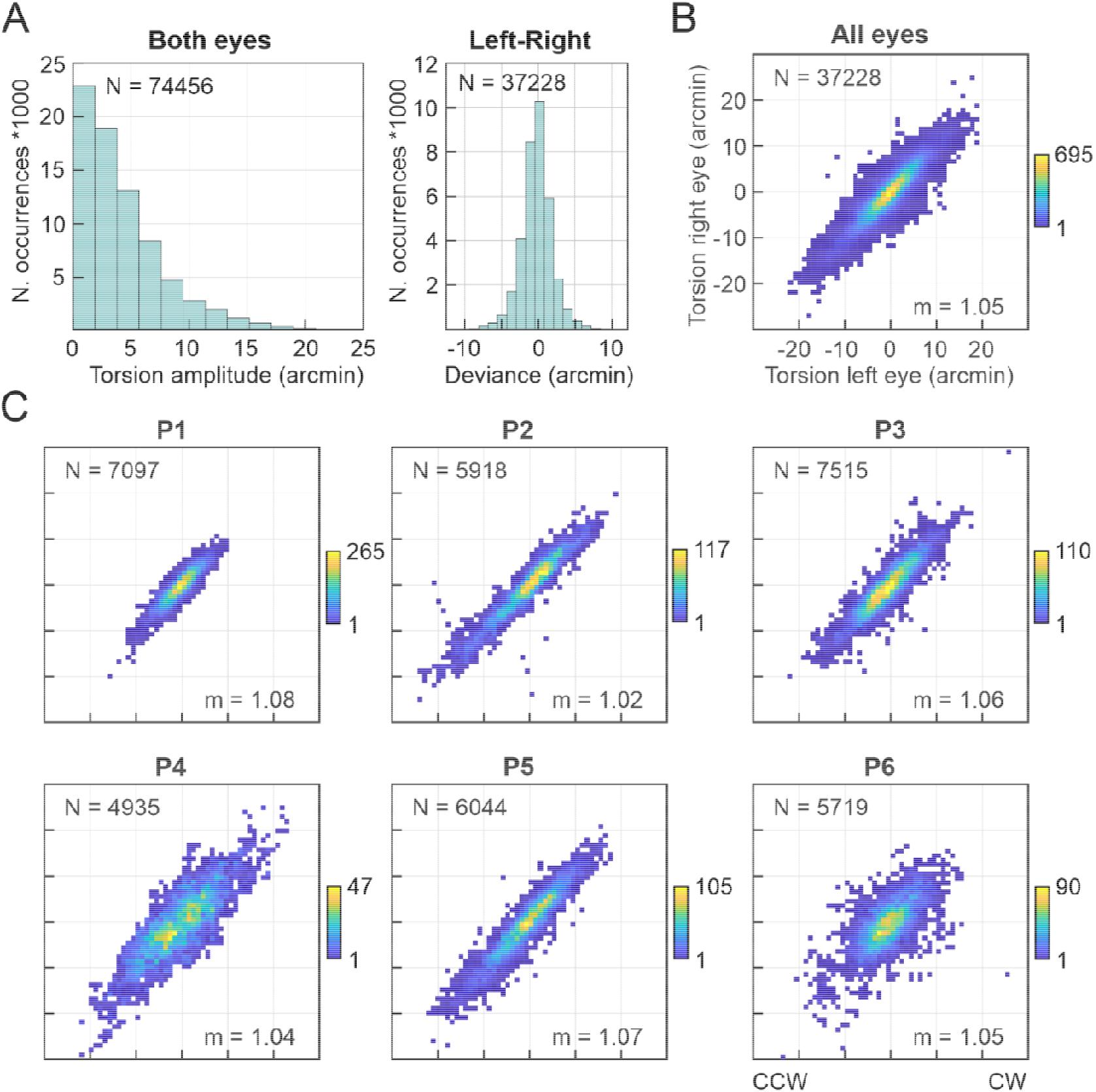
Coupling of cyclotorsion between fellow eyes. A: Histogram of the frame-wise torsion amplitudes of all eyes and participants and torsional amplitude deviance between all fellow eyes (computed as left-right). B: Correlation of torsional amplitudes across all eyes. Positive values represent clockwise, negative values represent counter-clockwise rotation. Data has been binned to 1 arcmin squares and are color coded for the number of occurrences in each bin. C: The same as in B, shown for each participant individually (P1-P6).

As a further observation, torsional motion corresponded to horizontal and vertical movement patterns of fixational eye movements, with gradual changes during drift and abrupt changes during saccades. The average torsional velocity change during a microsaccades was 2.32 arcmin/ms (SD: 1.57 arcmin/ms, Range: 0 - 7.56 arcmin/ms). Between microsaccades, i.e. during drift, the average torsional velocity change was 0.09 arcmin/ms (SD: 0.08 arcmin/ms, Range: 0 - 0.34 arcmin/ms).

## Discussion

We demonstrate an improved design of a split-field binocular scanning laser ophthalmoscope for high-resolution measurement of fixational eye movements. For validation, binocular fixational eye movements including cyclotorsion were assessed in six healthy participants.

The instrument described here offers some technical improvements to a similar split-field binocular SLO that was demonstrated earlier (Stevenson, Sheehy & Roorda, 2016). First, 3 × 3-degree imaging fields were used, allowing observation of larger eye movements. Fixational eye movements range from ∼11-60 arcsec (Tremor) to 7.5 ± 1.5 arcmin (Microsaccades) (Bowers, Boehm & Roorda, 2019; Montes, Bennett, Bensinger, Rani, Sherkat, Zhao & Sheehy, 2022), which would put our method in the position to cover most fixational situations while the larger imaging rasters can be used as a retinal display system for retina-contingent visual psychophysics (Yang, Arathorn, Tiruveedhula, Vogel & Roorda, 2010). Two independent Badal optometers allowed correction of ocular lower order aberrations, the largest factor reducing image quality in ophthalmic imaging (Steven, Sulai, Cheong, Bentley & Dubra, 2018). This will allow to increase observation numbers in normal participants and patients, given that eyes do not have to be preselected based on favorable refractive states. Interpupillary distance could be adjusted using motorized platforms, which together with an optional pupil monitor camera allowed relatively easy and quick binocular alignment. This is likely to increase imaging and workflow efficiency by decreasing chair time. Finally, binocular vergence angles could be independently induced by motorized rotation platforms, adding experimental options for binocular vision science experiments. Our system was built and optically optimized to accommodate a second light channel in the future for multi-wavelength binocular micro-stimulation (Harmening, Tuten, Roorda & Sincich, 2014).

Similar to other monocular SLO tracking systems (Sheehy, Yang, Arathorn, Tiruveedhula, Boer & Roorda, 2012), we could demonstrate high temporal and spatial resolution for both horizontal and vertical eye motion estimation in both eyes. Given the split-field design in our system, the right and left eye’s image content is recorded quasi-simultaneously, removing the need of temporal synchronization. In reality, the right and left signals are recorded block-wise (R, L, R, L, …), with a constant temporal offset between right and left signal onsets equal to half of the line rate (∼6.7 µs), which is less than 0.5% of the temporal sampling rate of about 1 ms, and thus negligible.

With SLO-based retinal tracking, Stevenson, Roorda & Kumar (2010) measured vertical vergence during steady fixation with a standard deviation of 1–2 arcmin. Horizontal vergence had more variability, ranging from 2 to 10 arcmin. We measured an average vergence motion of 1.18 arcmin (SD: 1.21 arcmin) in the horizontal, and 0.56 arcmin (SD: 0.53 arcmin) in the vertical direction. Bowers, Boehm & Roorda (2019) used an AOSLO and found the standard deviation of motion signals to be 5.10 ± 0.66 arcseconds horizontally and 5.51 ± 0.57 arcseconds vertically during steady fixation. Average microsaccade amplitude was measured at 7.5 ± 1.5 arcmin and average drift amplitude at 3.8 ± 0.9 arcmin. Our position signals had a standard deviation of 4.79/2.58 arcsec (horizontal/vertical), and average microsaccade amplitude of 13.46 arcmin and an average drift amplitude of 2.16 arcmin. Measurement performance was thus similar to invasive tracking. For instance, Riggs & Ratliff (1951) mounted mirrors on plastic contact lenses fitted directly to the moving eyeball. The system was able to record eye movements (horizontal, vertical, and torsional components) smaller than one arc minute. Dual Purkinje image tracker (Cornsweet & Crane, 1973; Crane et al., 1985) use a non-invasive optical method and achieve a precision of around 1 arcmin. For an excellent comparative overview of eye motion measurement precision across techniques see Sheehy et al. (2012).

While temporal and spatial resolution of SLO-based retinal tracking is equal or superior to commercial video based binocular eye trackers, they will only track motion amplitudes that are on the order of the imaging field size. Eye motion that produces image content with insufficient overlap to a common reference frame cannot be estimated reliably with such a system (Stevenson, Roorda & Kumar, 2010). This makes SLO-based retinal tracking ideal to study fixational eye motion, given their smaller amplitude (Rolfs, 2009). If image acquisition in a retinal imager is fast enough, temporally adjacent video frames can contain sufficient spatial overlap that removes the need of a common reference frame, and thus allows out-of-field tracking (Szkulmowski et al., 2020). Such an approach may offer, on the other hand, not enough spatial resolution to resolve retinal structure of interest, which may be important if gaze behavior and retinal cell topography is wished to be linked (Reiniger, Domdei, Holz & Harmening, 2021; Ratnam, Domdei, Harmening & Roorda, 2017; Harmening, Tuten, Roorda & Sincich, 2014)

Our data allowed analysis of the binocular coupling of FEM. With regard to microsaccades, it is widely accepted that they follow the same kinetics as other saccades (Zuber, Stark & Cook, 1965), establishing a microsaccade-saccade continuum that extends to free-viewing conditions (see, for example, Otero-Millan, Troncoso, Macknik, Serrano-Pedraza & Martinez-Conde, 2008). Whether microsaccades occur as a cyclopic phenomenon or could be generated monocularly is part of an ongoing debate in eye movement research: while early and recent studies using contact lens-based eye-tracking or binocular recordings from high-resolution search coil or Dual-Purkinje-image eye-tracking systems reported that microsaccades were highly conjugate between the two eyes (Krauskopf et al, 1960; Schulz, 1984; Fang, Gill, Poletti, & Rucci, 2018), several recent reports from video-based eye tracking studies showed and discussed the existence and prevalence of monocular microsaccades (see for example: Engbert & Kliegl, 2003; Martinez-Conde, Macknick, Troncoso, & Dyar, 2006; Gautier, Bedell, Siderov, & Waugh, 2016; but also: Kloke, Jaschinski, Jainta, 2009; Moller, Laursen, Tygesen & Sjolie, 2002 and Holmqvist & Blignaut, 2020 for methodological issues). Our data adds to this debate in favour for highly conjugate microsaccades: true monocular microsaccades were not present in our data.

In this work, cyclotorsional motion during fixation was extracted by analysis of the sawtooth pattern artifact in the horizontal movement track, as suggested in prior studies (Stevenson, Roorda & Kumar, 2010; Bowers, Boehm, & Roorda, 2019). After calibrating this linkage with image data that contained artificial rotation (see Methods), cyclotorsional movement could be estimated with an angular resolution of less than 1 arcmin. We note however that SLO-based torsion signals are theoretically confounded by the same artifacts as position signals are. High-speed motion path estimation from strip-wise image registration in a scanning system makes use of the fact that eye motion causes image distortions in each frame. This is because retinal structures are captured in a continuously updating (scanning) video frame as they move, and because the scanning speed is slower than the fastest occurring retinal motion. At the same time, such intra-frame image distortions pose a limit to creating a true, i.e. undistorted, representation of the unmoving retina. Because intra-frame distortions will be present in the image material used to construct a reference frame, their spatial signature will then show up in the motion path itself. Distortion-free reference frame generation is an ongoing topic in SLO-based eye motion research (Shenoy, Fong, Tan, Roorda & Ng, 2021; Bedggood & Metha, 2017; Bedggood & Metha, 2019), and accurate torsion estimation, like position estimation, will benefit from its success.

We found that torsional motion was, like microsaccades and drift, largely coupled between the two eyes, and, in accordance with earlier work, often occurred with or immediately after a saccade (Murdison, Blohm & Bremmer, 2019). Thus, our data tentatively suggests that torsional eye movements correct slight misallocations of the eyes after saccades (Howard, 2012) in a conjugate fashion. More data for different fixation stimuli is clearly needed to evaluate the functional role of such fixational eye movements in respect of single eye or binocular coordinated processes. The presented binocular scanning laser ophthalmoscope promises to be an apt setup for such research.

## Commercial relationships disclosure

**J Hofmann,** None; **L Domdei,** None; **S Jainta,** None; **W Harmening,** None.

## Acknowledgements

We thank Austin Roorda and Pavan Tiruveedhula for technical resources. This study was funded by a research grant of the Dr. Eberhard and Hilde Rüdiger Stiftung (BINOSLO), and by the Emmy Noether-Program of the German Research Foundation (DFG, Ha 5323/5-1,2).

## References

Barlow, H. B. (1952). Eye movements during fixation. The Journal of Physiology, 116(3), 290, https://doi.org/10.1113/jphysiol.1952.sp004706.

Bedggood, P., & Metha, A. (2017). De-warping of images and improved eye tracking for the scanning laser ophthalmoscope. PloS one, 12(4), e0174617, https://doi.org/10.1371/journal.pone.0174617.

Bedggood, P., & Metha, A. (2019). Mapping flow velocity in the human retinal capillary network with pixel intensity cross correlation. PloS one, 14(6), e0218918, https://doi.org/10.1371/journal.pone.0218918.

Bowers, N. R., Boehm, A. E., & Roorda, A. (2019). The effects of fixational tremor on the retinal image. Journal of vision, 19(11), 8–8, https://doi.org/10.1167/19.11.8.

Chung, S. T., Kumar, G., Li, R. W., & Levi, D. M. (2015). Characteristics of fixational eye movements in amblyopia: Limitations on fixation stability and acuity? Vision research, 114, 87–99, https://doi.org/10.1016/j.visres.2015.01.016.

Cornsweet, T. N., & Crane, H. D. (1973). Accurate two-dimensional eye tracker using first and fourth Purkinje images. JOSA, 63(8), 921–928, https://doi.org/10.1364/JOSA.63.000921.

Crane, H. D., & Steele, C. M. (1985). Generation-V dual-Purkinje-image eyetracker. Applied optics, 24(4), 527–537, https://doi.org/10.1364/AO.24.000527.

Engbert, R., & Kliegl, R. (2003). Microsaccades uncover the orientation of covert attention. Vision research, 43(9), 1035–1045, https://doi.org/10.1016/S0042-6989(03)00084-1.

Fang, Y., Gill, C., Poletti, M., & Rucci, M. (2018). Monocular microsaccades: Do they really occur?. Journal of vision, 18(3), 18–18, https://doi.org/10.1167/18.3.18.

Flitcroft, D. I., Judge, S. J., & Morley, J. W. (1992). Binocular interactions in accommodation control: effects of anisometropic stimuli. Journal of Neuroscience, 12(1), 188–203, https://doi.org/10.1523/JNEUROSCI.12-01-00188.1992.

Gautier, J., Bedell, H. E., Siderov, J., & Waugh, S. J. (2016). Monocular microsaccades are visual-task related. Journal of vision, 16(3), 37–37, https://doi.org/10.1167/16.3.37.

Gómez-Vieyra, A., Dubra, A., Malacara-Hernández, D., & Williams, D. R. (2009). First-order design of off-axis reflective ophthalmic adaptive optics systems using afocal telescopes. Optics express, 17(21), 18906–18919, https://doi.org/10.1364/OE.17.018906.

Harmening, W. M., Tuten, W. S., Roorda, A., & Sincich, L. C. (2014). Mapping the perceptual grain of the human retina. Journal of Neuroscience, 34(16), 5667–5677, https://doi.org/10.1523/JNEUROSCI.5191-13.2014.

Holmqvist, K., & Blignaut, P. (2020). Small eye movements cannot be reliably measured by video-based P-CR eye-trackers. Behavior research methods, 52(5), 2098–2121, https://doi.org/10.3758/s13428-020-01363-x.

Howard, I. P. (2012). Perceiving in depth, volume 1: basic mechanisms. Oxford University Press.

Kloke, W. B., Jaschinski, W., & Jainta, S. (2009). Microsaccades under monocular viewing conditions. Journal of Eye Movement Research, 3(1), https://doi.org/10.16910/jemr.3.1.2.

Ko, H. K., Poletti, M., & Rucci, M. (2010). Microsaccades precisely relocate gaze in a high visual acuity task. Nature neuroscience, 13(12), 1549–1553, https://doi.org/10.1038/nn.2663.

Krauskopf, J., Cornsweet, T. N., & Riggs, L. A. (1960). Analysis of eye movements during monocular and binocular fixation. JOSA, 50(6), 572–578, https://doi.org/10.1364/JOSA.50.000572.

Leigh, R. J., & Zee, D. S. (2006). The neurology of eye movements fourth edition. CONTEMPORARY NEUROLOGY SERIES, 70(1).

Martinez-Conde, S. (2006). Fixational eye movements in normal and pathological vision. Progress in brain research, 154, 151–176, https://doi.org/10.1016/S0079-6123(06)54008-7.

Martinez-Conde, S., Macknik, S. L., & Hubel, D. H. (2004). The role of fixational eye movements in visual perception. Nature reviews neuroscience, 5(3), 229–240, https://doi.org/10.1038/nrn1348.

Martinez-Conde, S., Otero-Millan, J., & Macknik, S. L. (2013). The impact of microsaccades on vision: towards a unified theory of saccadic function. Nature Reviews Neuroscience, 14(2), 83–96, https://doi.org/10.1038/nrn3405.

McCrea, R. A., Strassman, A., May, E., & Highstein, S. M. (1987). Anatomical and physiological characteristics of vestibular neurons mediating the horizontal vestibulo-ocular reflex of the squirrel monkey. Journal of Comparative Neurology, 264(4), 547–570, https://doi.org/10.1002/cne.902640408.

Møller, F., Laursen, M., Tygesen, J., & Sjølie, A. (2002). Binocular quantification and characterization of microsaccades. Graefe’s archive for clinical and experimental ophthalmology, 240(9), 765–770, https://doi.org/10.1007/S00417-002-0519-2.

Montes, S. Y. C., Bennett, D., Bensinger, E., Rani, L., Sherkat, Y., Zhao, C., & Sheehy, C. K. (2022). Characterizing Fixational Eye Motion Variance Over Time as Recorded by the Tracking Scanning Laser Ophthalmoscope. Translational Vision Science & Technology, 11(2), 35–35, https://doi.org/10.1167/tvst.11.2.35.

Murdison, T. S., Blohm, G., & Bremmer, F. (2019). Saccade-induced changes in ocular torsion reveal predictive orientation perception. Journal of Vision, 19(11), 10–10, https://doi.org/10.1167/19.11.10.

Murray, J., Gupta, P., Dulaney, C., Garg, K., Shaikh, A. G., & Ghasia, F. F. (2022). Effect of Viewing Conditions on Fixation Eye Movements and Eye Alignment in Amblyopia. Investigative Ophthalmology & Visual Science, 63(2), 33–33, https://doi.org/10.1167/iovs.63.2.33.

Otero-Millan, J., Macknik, S. L., Langston, R. E., & Martinez-Conde, S. (2013). An oculomotor continuum from exploration to fixation. Proceedings of the National Academy of Sciences, 110(15), 6175–6180, https://doi.org/10.1073/pnas.1222715110.

Otero-Millan, J., Macknik, S. L., & Martinez-Conde, S. (2014). Fixational eye movements and binocular vision. Frontiers in integrative neuroscience, 8, 52, https://doi.org/10.3389/fnint.2014.00052.

Otero-Millan, J., Troncoso, X. G., Macknik, S. L., Serrano-Pedraza, I., & Martinez-Conde, S. (2008). Saccades and microsaccades during visual fixation, exploration, and search: foundations for a common saccadic generator. Journal of vision, 8(14), 21–21, https://doi.org/10.1167/8.14.21.

Poletti, M., Listorti, C., & Rucci, M. (2010). Stability of the visual world during eye drift. Journal of Neuroscience, 30(33), 11143–11150, https://doi.org/10.1523/JNEUROSCI.1925-10.2010.

Poletti, M., & Rucci, M. (2015). A compact field guide to the study of microsaccades: Challenges and functions. Vision research, 118, 83–97, https://doi.org/10.1016/j.visres.2015.01.018.

Ratnam, K., Domdei, N., Harmening, W. M., & Roorda, A. (2017). Benefits of retinal image motion at the limits of spatial vision. Journal of vision, 17(1), 30–30, https://doi.org/10.1167/17.1.30.

Reiniger, J. L., Domdei, N., Holz, F. G., & Harmening, W. M. (2021). Human gaze is systematically offset from the center of cone topography. Current Biology, 31(18), 4188–4193, https://doi.org/10.1016/j.cub.2021.07.005.

Riggs, L. A., & Ratliff, F. (1951). Visual acuity and the normal tremor of the eyes. Science, 114(2949), 17–18, https://doi.org/10.1126/science.114.2949.17.

Rolfs, M. (2009). Microsaccades: small steps on a long way. Vision research, 49(20), 2415–2441, https://doi.org/10.1016/j.visres.2009.08.010.

Rucci, M., & Victor, J. D. (2015). The unsteady eye: an information-processing stage, not a bug. Trends in neurosciences, 38(4), 195–206, https://doi.org/10.1016/j.tins.2015.01.005.

Schor, C. M., & Ciuffreda, K. J. (1983). Accommodative Vergence and Accommodation in Normals, Amblyopes, and Strabismics. Vergence Eye Movements: Basic, 143.

Schulz, E. (1984). Binocular micromovements in normal persons. Graefe’s archive for clinical and experimental ophthalmology, 222(2), 95–100.

Sheehy, C. K., Yang, Q., Arathorn, D. W., Tiruveedhula, P., de Boer, J. F., & Roorda, A. (2012). High-speed, image-based eye tracking with a scanning laser ophthalmoscope. Biomedical optics express, 3(10), 2611–2622, https://doi.org/10.1364/BOE.3.002611.

Shenoy, J., Fong, J., Tan, J., Roorda, A., & Ng, R. (2021). R-SLAM: Optimizing Eye Tracking from Rolling Shutter Video of the Retina. In Proceedings of the IEEE/CVF International Conference on Computer Vision (pp. 4852–4861).

Simon, F., Schulz, E., Rassow, B., & Haase, W. (1984). Binocular micromovement recording of human eyes: - methods. Graefe’s archive for clinical and experimental ophthalmology, 221(6), 293–298., https://doi.org/10.1007/BF02134127.

Spauschus, A., Marsden, J., Halliday, D. M., Rosenberg, J. R., & Brown, P. (1999). The origin of ocular microtremor in man. Experimental Brain Research, 126(4), 556–562, https://doi.org/10.1007/s002210050764.

Steinman, S. B., Steinman, B. A., & Garzia, R. P. (2000). Foundations of binocular vision: a clinical perspective. McGraw-Hill Education/Medical.

Steven, S., Sulai, Y. N., Cheong, S. K., Bentley, J., & Dubra, A. (2018). Long eye relief fundus camera and fixation target with partial correction of ocular longitudinal chromatic aberration. Biomedical Optics Express, 9(12), 6017–6037, https://doi.org/10.1364/BOE.9.006017.

Stevenson, S. B., Roorda, A., & Kumar, G. (2010). Eye tracking with the adaptive optics scanning laser ophthalmoscope. In Proceedings of the 2010 symposium on eye-tracking research & applications (pp. 195–198), https://doi.org/10.1145/1743666.1743714.

Stevenson, S. B., Sheehy, C. K., & Roorda, A. (2016). Binocular eye tracking with the tracking scanning laser ophthalmoscope. Vision research, 118, 98–104, https://doi.org/10.1016/j.visres.2015.01.019.

Szkulmowski, M., Meina, M., Bartuzel, M., Wrobel, K., Tamborski, S., Nowakowski, M. & Szkulmowska, A. (2020). Ultrafast retinal tracker for quantification of fixational and saccadic motion of the human eye. Investigative Ophthalmology & Visual Science, 61(7), 1849–1849.

Tweed, D. (1997). Visual-motor optimization in binocular control. Vision research, 37(14), 1939–1951, https://doi.org/10.1016/S0042-6989(97)00002-3.

Van Rijn, L. J., Van der Steen, J., & Collewijn, H. (1992). Visually induced cycloversion and cyclovergence. Vision Research, 32(10), 1875–1883, https://doi.org/10.1016/0042-6989(92)90048-N.

Van Rijn, L. J., Van der Steen, J., & Collewijn, H. (1994). Instability of ocular torsion during fixation: cyclovergence is more stable than cycloversion. Vision Research, 34(8), 1077–1087, https://doi.org/10.1016/0042-6989(94)90011-6.

Vogel, C. R., Arathorn, D. W., Roorda, A., & Parker, A. (2006). Retinal motion estimation in adaptive optics scanning laser ophthalmoscopy. Optics express, 14(2), 487–497, https://doi.org/10.1364/OPEX.14.000487.

Yang, Q., Arathorn, D. W., Tiruveedhula, P., Vogel, C. R., & Roorda, A. (2010). Design of an integrated hardware interface for AOSLO image capture and cone-targeted stimulus delivery. Optics express, 18(17), 17841–17858, https://doi.org/10.1364/OE.18.017841.

Zhou, W., & King, W. M. (1998). Premotor commands encode monocular eye movements. Nature, 393(6686), 692–695, https://doi.org/10.1038/31489.

Zuber, B. L., Stark, L., & Cook, G. (1965). Microsaccades and the velocity-amplitude relationship for saccadic eye movements. Science, 150(3702), 1459–1460, https://doi.org/10.1126/science.150.3702.1459.

